# A Seven-Protein Assembly Promotes Stability, Neutralization and Secretion of the T7SSb LXG-effector TelE

**DOI:** 10.64898/2026.03.12.711312

**Authors:** Palak Agrawal, Antonina Gonet, Doreen Toko, Johanna Gorkisch, Dorothée Raoux-Barbot, Laurence du Merle, Eleonore Bouscasse, Mariette Matondo, Ariel Mechaly, Wooi Keong Teh, Armel Bezault, Alexandre Chenal, Marco Bellinzoni, Shaynoor Dramsi, Francesca Gubellini

## Abstract

*Streptococcus gallolyticus* subsp. *gallolyticus (SGG)* is a gut pathobiont associated with colorectal cancer. Like many Firmicutes, *SGG* utilizes a specialized Type VII Secretion System (T7SSb) to export WXG100 and LXG proteins, the latter involved in bacterial competition. We previously identified TelE, an LXG protein whose C-terminus mediates membrane pore formation in *Escherichia coli*. In *SGG* UCN34, TelE (Gallo_0562) is co-expressed with six proteins (Gallo_0559 to Gallo_0565), including its immunity protein TipE (Gallo_0565). Here we show that the absence of those co-expressed proteins affects TelE stability and secretion. These proteins associate with TelE to form a soluble and stable seven-protein complex. Gallo_0559 and Gallo_0560 interact with the N-terminal LXG domain of TelE, Gallo_0561 binds to its central region, while the six-transmembrane protein Gallo_0563, together with Gallo_0564 and TipE associates with its C-terminal domain. These findings describe a new modular complex that stabilizes TelE while reducing its toxicity and optimizing its T7SSb-mediated delivery.

## INTRODUCTION

The Type VII secretion system in Bacillota (T7SSb) mediates interbacterial competition through the secretion of diverse substrates, including WXG100 proteins and toxic LXG effectors (Abdallah et al. 2007; Spencer and Doran 2022; Boardman et al. 2023; Garrett and Palmer 2024). The LXG effectors target rival bacteria and can also modulate the host environment (Tran et al. 2021; Bowman and Palmer 2021, Boardman et al. 2023). These proteins typically feature a conserved N-terminal LXG domain (about 200 residues; PF04740/IPR006829) containing a highly conserved region, and a variable C-terminal domain conferring the effector’s specific function. These toxic activities include NADase, lipid II phosphatase, ADP ribosyl transferase, nucleases, and membrane depolarization. Bacteria such as *Bacillus subtilis*, *Staphylococcus aureus*, *Streptococcus intermedius*, and *Enterococcus faecalis* secrete these effectors to gain a survival advantage within competitive microbial environment (Holberger et al. 2012; Cao et al. 2016; Ohr et al. 2017; Ulhuq et al. 2020; Alhajjar et al. 2020; Chatterjee et al. 2021; Tassinari et al. 2022; Boardman et al. 2023; Garrett and Palmer 2024).

LXG effectors are typically encoded within the so-called ‘LXG operon’. Frequently, genes located upstream of the LXG effectors encode a novel class of LXG-associated α-helical proteins, named Lap, which are required for effector secretion (Klein et al. 2022, 2024; Yang et al. 2023). Lap proteins contain various conserved domains of unknown function in their sequence, including DUF3130 in Lap1, DUF3958 in Lap2, DUF5344 in Lap3 and DUF5082 in Lap4 (Klein et al. 2024). In *S. intermedius*, Lap proteins form trimeric complexes with the LXG domains of the effectors, likely stabilizing them and targeting them to the T7SSb machinery for export (Klein et al. 2024). In Group B Streptococcus, the expression of at least one Lap protein (BlpA1 or BlpA2) is necessary and sufficient for the stability and toxicity of the LXG effector BltA (Job et al. 2024).

Additional analyses of LXG operons have uncovered proteins containing DUF4176, which were proposed to function as specific molecular chaperones stabilizing the cognate LXG effector (Gkragkopoulou et al. 2025; Wu et al. 2025). LXG operons also encode the cognate immunity proteins. Moreover, a recent study described two distinct families of immunity proteins mediating compartment-specific neutralization of the membrane-depolarizing LXG effector EsxX in *S. aureus*. Notably, a novel class of multi-spanning membrane proteins was shown protecting the producing cells from the extracellular toxicity of the effector (Alcock et al. 2025).

Collectively, these studies suggest that LXG effectors bind to multiple proteins required for their structural stabilization, targeting to their T7SSb machinery, and neutralization of their toxic activity. Nevertheless, key questions remain regarding how these processes are coordinated and integrated with the LXG effectors secretion through the T7SSb pathway.

*Streptococcus gallolyticus* subsp. *gallolyticus* (*SGG*), our model bacterium, was the first intestinal bacterium associated with colorectal cancer (McCOY and Mason 1951; Abdulamir et al. 2011; Boleij et al. 2011; Pasquereau-Kotula et al. 2018). Microbiota dysbiosis is now recognized as a critical factor in colon tumor development (White and Sears 2024). Deletion of the entire T7SSb secretion system in *SGG* strain TX20005 significantly reduced bacterial adherence to host colonic cells and hampered the subsequent tumor development (Taylor et al. 2021). We previously characterized the LXG-effector TelE in *SGG* strain UCN34 (Teh et al. 2023). The gene encoding TelE (*gallo_0562*) is located downstream the *ess* locus encoding the T7SSb machinery, as part of a seven-gene operon (here called ‘LXG_TelE_ operon’) (Teh et al. 2023). Three genes are found upstream of TelE. Proteins encoded by the first two genes, *gallo_0559* and *gallo_0560*, co-secreted with TelE, are alpha-helical proteins homologous to Lap proteins (Klein et al. 2022, 2024, Yang et al., 2023). The third gene, *gallo_0561,* encodes a DUF4176 protein, same as the Lcp <u>c</u>haperones reported in (Gkragkopoulou et al. 2025; Wu et al. 2025). The remaining three proteins encoded by the genes downstream of *telE* are the immunity protein TipE (Gallo_0565) and its structural homolog Gallo_0564 (both DUF5085 family members), and a six-transmembrane protein Gallo_0563 without a predicted DUF domain (Blum et al. 2025; Paysan-Lafosse et al. 2025).

In this study, we characterized a soluble, high-molecular weight complex formed by TelE and its six co-expressed proteins. Our findings demonstrate that these TelE-associated proteins function collaboratively to stabilize the protein complex for TelE secretion and its intracellular toxicity neutralization. Two of the six associated proteins were co-secreted alongside TelE via the T7SSb machinery. Based on our crystallography and cryo-electron microscopy (cryo-EM) analyses, we propose a model in which these proteins assemble in a modular complex within the cytosolic compartment. This complex transiently inactivates the effector’s toxicity, while maintaining it in a secretion-competent state.

## RESULTS

### Characterization of the genes co-expressed with *telE* and *tipE*

To identify the genes involved in the homeostasis and/or secretion of TelE, we generated in-frame deletion mutants for each gene in the LXG_TelE_ operon *gallo_0559* to *gallo_0565* in the *SGG* strain UCN34 (Δ*0559*, Δ*0560*, Δ*0561*, Δ*0562*, Δ*0563*, Δ*0564*) (**Figure 1A**). We also constructed a mutant deleted for the entire LXG_TelE_-operon (Δ*0559-0565*). The Δ*essC* mutant constructed previously (Teh et al., 2023), which is unable to express T7SSb’s motor ATPase, was used as the T7SSb-defective control. Under our laboratory conditions, the mutants are not different than the UCN34 wild-type strain in growth (**Figure S1A**). To rule out polar effects, we measured *telE* transcript levels in all mutants using quantitative RT-PCR (**Figure S1B**). As expected, *telE* transcripts were absent in the Δ*0562* and Δ*0559-0565* mutants, validating the primers used. TelE transcript levels were similar in all other mutants, including Δ*0559*, Δ*0560*, Δ*0561* as the WT strain, confirming the absence of polar effects.

**Figure 1.**
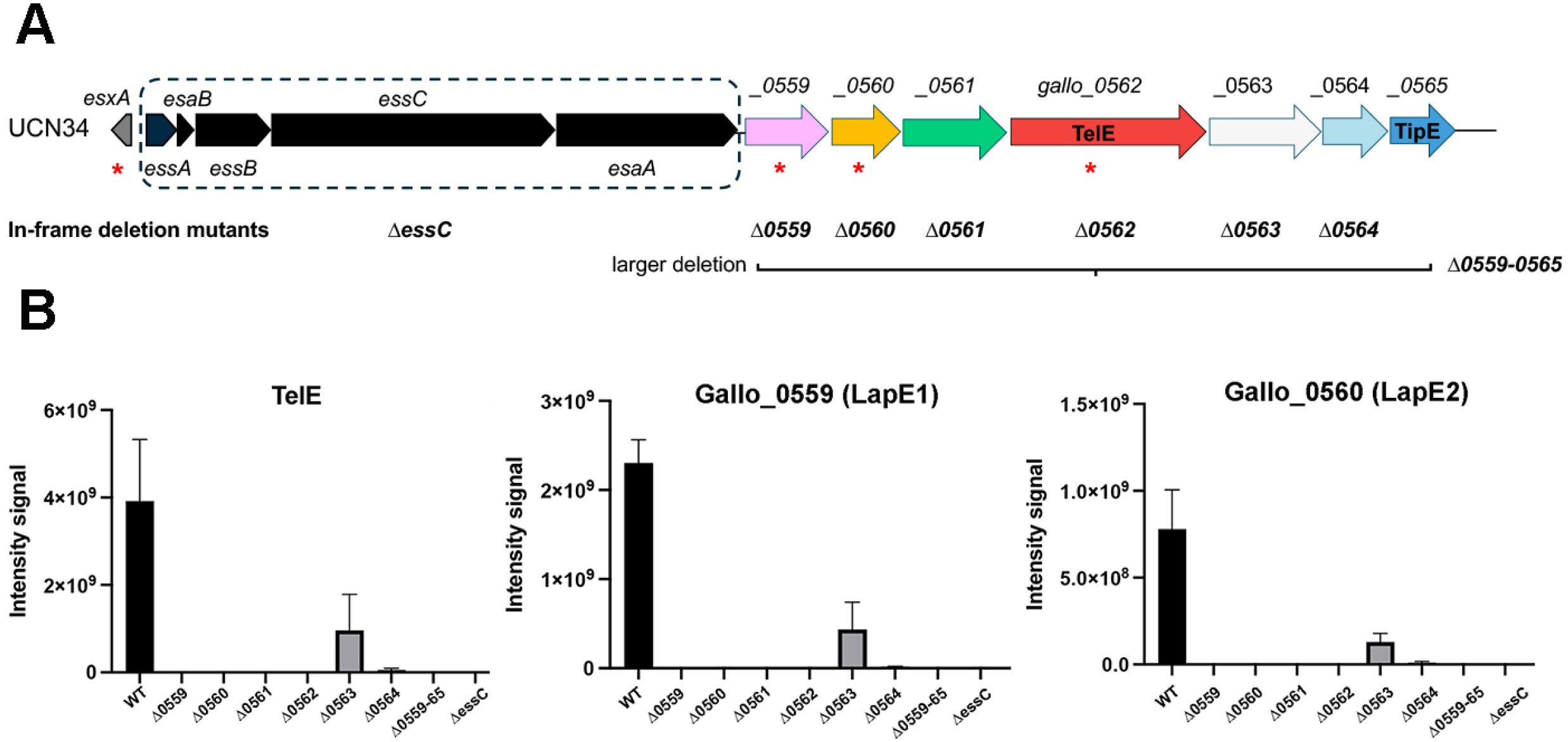
All the six genes co-expressed with *telE* are necessary for TelE detection in the extracellular compartment. (A) Schematic of T7SSb locus in *SGG* UCN34 wild-type (WT) strain. The highly conserved core genes encoding the T7SSb machinery are shown in black within a dashed box whereas the downstream genes interacting with the LXG effector TelE are shown in various colors. The red asterisk highlights the proteins previously shown as secreted through the T7SSb machinery. (B) Mass spectrometry analysis of TelE, 59 and 60 protein abundances in the secretomes of UCN34 WT strain and single-gene deletion mutants *Δ0559*, *Δ0560*, *Δ0561*, *Δ0562* (telE), *Δ0563*, *Δ0564*, multi-gene deletion mutant (Δ*0559-65*) and T7SSb machinery defective Δ*essC* strain.

To assess the impact of these deletions on protein secretion, we compared the secretomes of the wild-type (WT) and mutant strains using mass spectrometry. Among the proteins identified (**Table S1**), we focused on the three proteins encoded within the LXG_TelE_ operon that were previously shown to be secreted by the T7SSb machinery in *SGG* UCN34 (Teh et al., 2023). These were: Gallo_0559 and Gallo_0560 (hereby shortened ‘59’ and ‘60’), and the LXG-effector TelE. Notably, deletion of any gene within the LXG_TelE_ operon strongly reduced the abundance of TelE, LapE1 and LapE2 in the secretome (**Figure 1B**), revealing the key role of these proteins in TelE stability and/or secretion.

### TelE and its binding partners

The TelE protein (514 residues total) contains an N-terminal LXG domain (residues 1-195) and a C-terminal glycine zipper (residues 458-480) that is critical for its intracellular membrane-permeabilizing activity in *E. coli* (Teh et al. 2023). Since TelE is a secreted protein, we investigated whether *E. coli* could be intoxicated by extracellularly administered TelE. To address this question, TelE was expressed in *E. coli*. Due to the inherent toxicity and instability of TelE in *E. coli* (Teh et al. 2023), we fused the Maltose Binding Protein (MBP) to the C-terminus of TelE, which significantly improved its stability and solubility. However, following MBP-tag cleavage, the untagged TelE remained intact only for a limited time (maximum 48 hours at 4°C) (**Figure 2A**).

**Figure 2.**
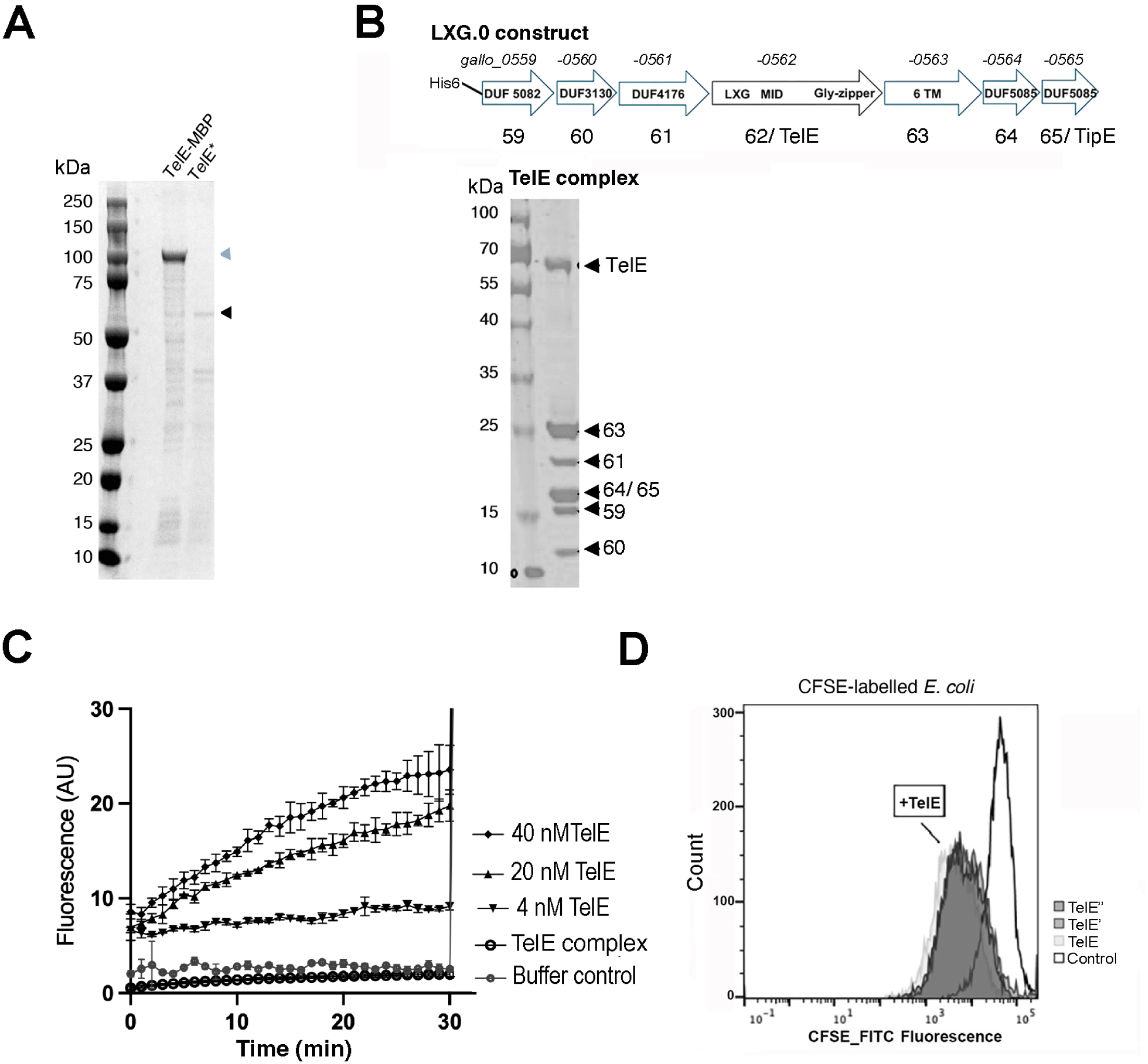
TelE, but not TelE-complex, permeabilizes lipid vesicles and bacterial membranes. (A) SDS-PAGE analysis of the purified TelE-MBP and untagged TelE proteins, resolved on 4-20% polyacrylamide gradient gels stained with Coomassie. (B) Top: schematic representation of the LXG.0 construct for co-expression and purification of the TelE-complex in *E. coli*. The size of each subunit in amino acids is as follows: 138 (59), 93 (60), 220 (61), 514 (TelE), 244 (63), 152 (64) and 151 (TipE). Their conserved features (such as DUFs and LXG domains) are indicated. MID indicates middle region of TelE. 6 TM= six transmembrane domains in the subunit 63. Bottom: SDS-PAGE analysis of the purified TelE-complex, resolved on 4-20% polyacrylamide gradient gel stained with Coomassie. (C) Changes in the fluorescence signals corresponding to the release of ANTS fluorescence upon LUV permeabilization (excitation: 360 nm, emission: 515 nm) induced by addition of purified TelE at various concentrations and of TelE-complex at 0.5 μM (≈ 40 nM TelE). At the end of the experiment, Triton X-100 was added as positive control. AU, arbitrary unit. (D) Flow cytometry analysis of CFSE-labeled *E. coli* cells incubated with purified TelE protein or with buffer control.

We reckoned that TelE could be stabilized by the proteins encoded in the LXG_TelE_ operon. Therefore, we expressed the entire LXG_TelE_ operon in *E. coli* (construct LXG.0, spanning from His6.*gallo_0559* to *gallo_0565*). Affinity purification yielded a multiprotein assembly (**Figure 2B**) hereafter referred to as the ‘TelE-complex’, composed of seven subunits, as confirmed by mass spectrometry (**Figure S2B**). Interestingly, the six-transmembrane protein Gallo_0563 co-purified with this soluble complex. For simplicity, Gallo_0561, Gallo_0563, and Gallo_0564 are hereafter referred to using their last two digits (proteins 61, 63 and 64).

Mass photometry analysis revealed that the purified TelE-complex had a mass of 202 kDa, with minor peaks at 141 kDa and 52 kDa (**Figure S2C**). To assess its stability, the complex was concentrated through a 100 kDa molecular cut-off membrane. No proteins were detected in the flow-through after filtration. In addition, Western blot analysis of both the concentrated and non-concentrated sample, using antibodies raised against the TelE complex, showed identical bands pattern, confirming the stability of the complex (**Figure S2D**).

Next, we tested the pore-forming activity of TelE and the TelE-complex in Large Unilamellar Vesicles (LUV), mimicking prokaryotic membranes composition (Murzyn et al. 2005). LUV were loaded with a fluorescence marker, 8-aminonaphthalene-1,3,6-trisulfonic acid (ANTS) and its quencher, p-xylene-bis-pyridinium bromide (DPX) at high concentration so that no fluorescence is released when they are both encapsulated in the LUVs. Upon pore formation in the liposomal membranes, ANTS and DPX are released into the medium, leading to fluorescence recovery of ANTS. Untagged TelE permeabilized the LUVs in a dose-dependent manner, whereas the TelE-complex displayed minimal activity, consistent with the presence of the immunity protein TipE (**Figure 2C**).

We then assessed TelE activity on living bacteria labelled with carboxyfluorescein succinimidyl ester (CFSE). CFSE passively diffusing in bacterial cells, being converted to its fluorescent form when inside the cytosol. *E. coli* cells labelled with the intracellular fluorescent dye CFSE were treated with purified TelE (**Figure 2D**). Within 10 min at 37 °C, TelE induced rapid dye leakage, comparable to the effect of the detergents Tween20 and Triton X100 used as positive control (**Figure S2A**, left panel). Control buffer containing β-DDM (N-dodecyl-β-D-maltoside) had negligible impact on bacterial permeabilization (**Figure S2A**, right panel).

### Structural analysis of the TelE-complex

To elucidate the molecular basis underlying the inter-subunits interactions within the TelE-complex, we employed X-ray crystallography and cryo-EM approaches. Given that the full TelE-complex resisted crystallization, we conducted limited proteolysis to generate sub-complexes suitable for this technique. The resulting proteolytic products were separated by size-exclusion chromatography, resolving as three distinct peaks (**Figure S3**). Among them, only peak #3 yielded crystals that diffracted to a resolution of 1.8 Å. Structural refinement revealed two trimeric assemblies arranged in a head-to-tail orientation within the asymmetric unit (**Figure S4A**). Each trimer included one copy of TelE (from residue 1 to 215, called TelE_1-215_) associated with proteins 59 and 60. Based on their structural homology with previously described Lap proteins (Klein et al. 2022, 2024, Yang et al., 2023), these proteins were renamed LapE1 and LapE2 respectively (**Figure 3A**, **Table S2**). Together, these three proteins assemble into an elongated structure, hereafter referred to as the ‘stalk’, with an overall length of 152 Å. The stalk exhibits variable thickness, measuring 32 Å at the interface where TelE interacts with LapE2 and narrowing down to 20 Å at its center (from Ser88 to Asp97 of TelE) (**Figure 3A**). This central region includes the N-terminal domain of TelE, which remained intact despite the proteolytic treatment. The first four residues (Met1 to Lys4) are buried in two antiparallel β-strands that are stabilized by 12 hydrogen bonds (**Figure 3A and S4B**). Multiple Sequence Alignment showed that these antiparallel β-strands have a certain degree of conservation, suggesting that this structural feature may have a functional role (**Figure S4C**). Each β-strand interacts with one Lap protein, creating a steric barrier blocking their interaction. LapE1 and LapE2 do not interact directly, each of them contacting exclusively the LXG domain of TelE. The interfaces between LapE1 and TelE (ΔG= −34.7 kcal/mol), and between TelE and LapE2 (ΔG= −36.9 kcal/mol), are stabilized by hydrophobic residues (**Figure S4E**). In addition to the ‘LXG domain’ encompassed within TelE_1-215_, two conserved sequences (YxxxD and NKxxDD) were observed on on LapE2 (YTKTD and NKSDDD) nearby the LxG motif on TelE (LQG) (**Figures 3A** and **S4D**).

**Figure 3:**
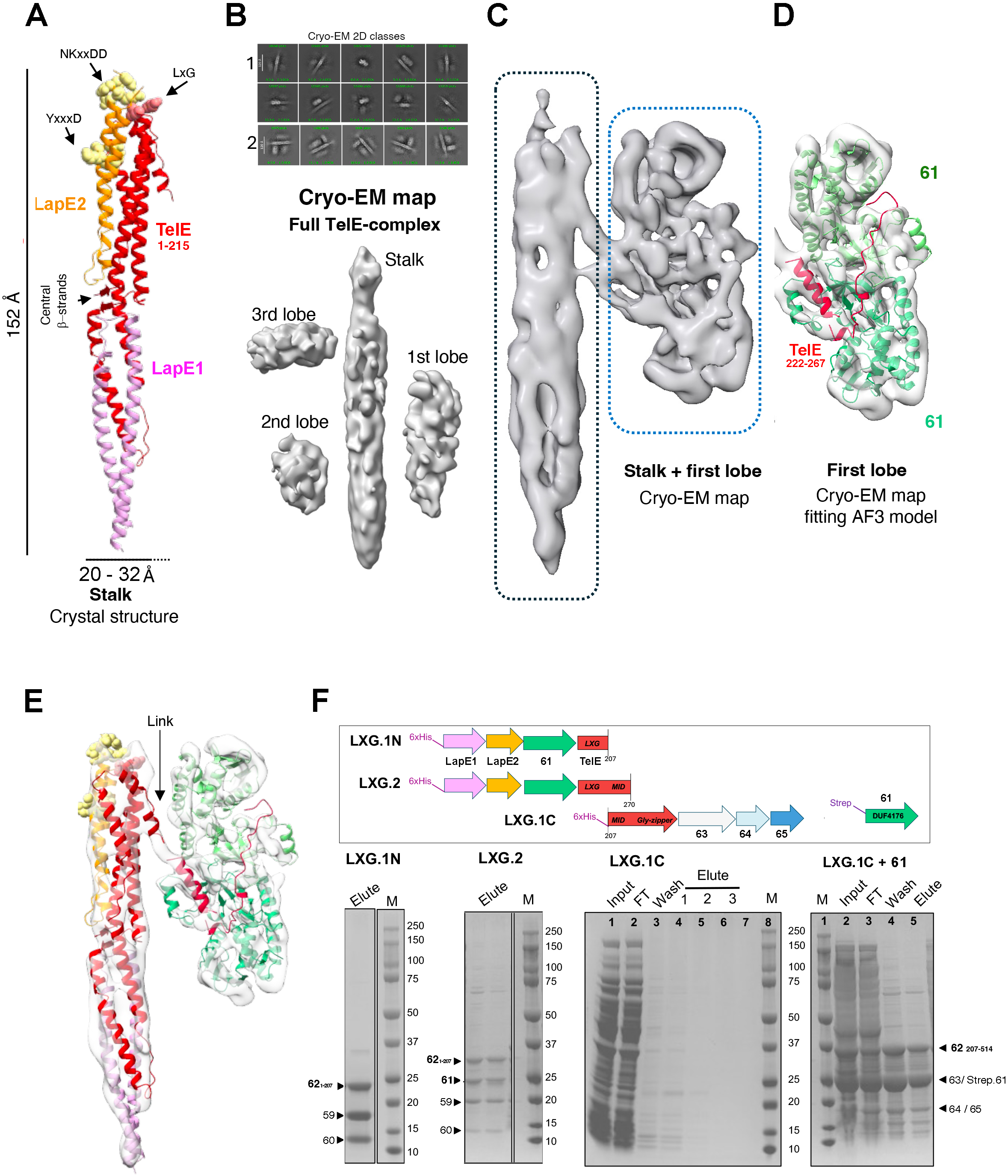
Structural features of the TelE-complex: Stalk and Lobe 1. (A) Crystallographic model of the trimer formed by subunits 59 - 60 and TelE (fragment 1-215) depicted in lavender, orange and red, respectively. (B) Representative cryo-EM 2D classes of the TelE complex, showing particles heterogeneity (panel 1). In panel 2, 2D classes displaying central stalk surrounded by three lobes. The corresponding cryo-EM map is shown below. (C) Cryo-EM reconstruction of the stalk and lobe 1, contoured at a threshold of 0.044 on ChimeraX. The stalk is indicated in dashed black box and lobe 1 in dashed blue box. (D) The AF3-predicted model of 61 dimer bound to TelE_222-267_ was fit into lobe 1 cryo-EM map (cross-correlation: 0.84). (E) The crystal structure of the TelE _1-215_-LapE1 (59)-LapE2(60) trimer together with AF3 model shown in D were fitted into the cryo-EM map of the stalk and lobe 1. (F) Top, Schematic representation of LXG.1N, LXG.2, LXG.1C and *strep-gallo_0561* constructs. Bottom, purified sub-complexes: LXG.1N, LXG.2 and LXG.1C co-expressed in the presence and absence of Strep-61 were analyzed by SDS-PAGE (4-20%) stained by Coomassie.

In parallel, we investigated the structure of the TelE-complex using cryo-EM. Analysis of particle images grouped into representative views (2D classes) revealed significant structural heterogeneity within the TelE-complex (**Figure 3B**, panel 1). The largest particles displayed an elongated structure surrounded by densities of globular domains (2D classes in **Figure 3B**, panel 2). Computational analysis of these particles produced a three-dimensional (3D) electron density map featuring a central stalk surrounded by three densities, hereafter designated as “first”, “second” and “third” lobe (**Figures 3B** bottom, and **S5A**). The resolution of this map was limited to a nominal 6.5 Å, likely due to the dynamic nature of the lobes, as evidenced by the differences in their relative orientations in the 2D classes (**Figure 3B**, panel 2).

To gain further insights into TelE complex, we analyzed the particles of the first and the second lobes separately. Particles presenting the stalk linked to the first lobe produced well-defined 2D classes (**Figure S4F,** panel 1). Their tree-dimensional reconstruction reached a nominal resolution of 4.5 Å (**Figures 3C** and **S4G**), despite limitations caused by preferential particle orientation. To interpret the cryo-EM map, we first fitted the crystal structure into the stalk density. In the first lobe, we fitted an AlphaFold (AF3) model of a subcomplex consisting of a dimer of subunit 61 interacting with 46 residues of TelE (Gly222 to Ile267) (**Figures 3D**, **3E and S4H**). By integrating the crystal structure with the model predicted by AF3, we propose that the α-helical region of TelE (Lys198 and Leu237) links the stalk and the first lobe (**Figure 3E**).

Consistent with this model, biochemical analysis showed that subunit 61 does not bind to a TelE construct comprising only the first 207 residues (expression of construct LXG.1N, **Figure 3F** and **Figure S6A**). In this context, only LapE1 and LapE2 remained bound to the TelE_1-207_ LXG domain, forming a stable trimeric complex. The absence of subunit 61 from this sub-complex was confirmed by SDS-PAGE and mass spectrometry (**Figures 3F and S6D**). In contrast, purification of the LXG.2 construct, which included the first 270 residues of TelE, yielded a complex that retained subunit 61, as verified by SDS-PAGE (**Figures 3F** and **S6B**) and mass spectrometry (**Figure S6D**).

We next investigated whether subunit 61 is required to stabilize the C-terminal region of TelE. Initial attempts to express the middle and C-terminal regions of TelE (residues 207-514) along with subunits 63, 64 and 65 (construct LXG.1C) were unsuccessful, as none of these proteins was detected in either the soluble or in the membrane fractions (**Figure 3F**). This suggests that the C-terminal region of TelE is inherently unstable. However, co-expression of the LXG.1C construct with Strep.61 subunit in *E. coli,* followed by affinity purification, yielded a soluble complex containing the C-terminus of TelE, along with subunits 63, 64, 65, and 61 (**Figures 3F** last gel on the right, and **S6C**). These results establish that Gallo_0561, a DUF4176 family protein, is essential for stabilizing the C-terminal region of TelE and for enabling the assembly of the full multiprotein complex.

We then analyzed the particles presenting the “second lobe” of the TelE-complex, which could be found in two forms depending on its position: either proximal or distal from the stalk (**Figure S4F,** panel 2). 2D classes of particles in the distal position were better defined **(Figure S4F** panel 3), enabling the generation of a 3D map at 4.2 Å resolution (**Figures 4A** and **S5B**). Analysis of the particles in proximal position (2D classes shown in **Figure S4F** panel 4) produced a comparable model, albeit at lower resolution (5.2 Å, **Figures 4B** and **S5C**). Interpretation of the distal and the proximal maps of the second lobe using AF3 indicated that it is primarily composed of subunits 64 and 65 (**Figure S5D**). To further understand how TelE interacts with these subunits, we generated an AF3 model of the full TelE-complex (**Figure S5E**), from which we extracted the region containing the 64–65 heterodimer along with the TelE segment that contacts it. This ‘cropped’ model was then fitted into the second lobe’s maps (distal and proximal) using ChimeraX (**Figures 4C and 4D**). In both maps, TelE interacts with the subunit 64 at two discrete sites: one near residue 344 (residues 345–349 *vs* 344–351) and another at the hinge between subunits 64 and 65 (residues 380–383 *vs* 384–387) (**Figures 4C and 4D**). In contrast, the interaction between TelE and subunit 65 (TipE) differs between the distal and proximal maps, with an empty groove observed exclusively in the distal map (**Figures 4C** *vs* **4D**). Here, the TelE residues 391–397 occupy a small density adjacent to the empty groove of TipE (**Figure 4C**), whereas in the proximal map, two α-helices fit the additional densities present on the back and within the groove of TipE (residues 388–397 and 401–421). As an alternative interpretation, this last density could accommodate the Glycine-rich region of TelE (residues 454–487, **Figure S5F**), as proposed for the TelE homolog EsxX (Alcock et al. 2025). However, in the AF3 models of the TelE complex, the glycine-rich region of TelE (responsible for pore formation) is always bound to the membrane subunit 63, rather than to TipE. We therefore propose that TipE may neutralize TelE toxicity by blocking its C-terminal region, upstream of the glycine-rich portion. The limited resolution of the available maps (including those in Alcock et al., 2025) does not allow a clear assignment of the interactions between TelE and the 64–65 dimer.

**Figure 4:**
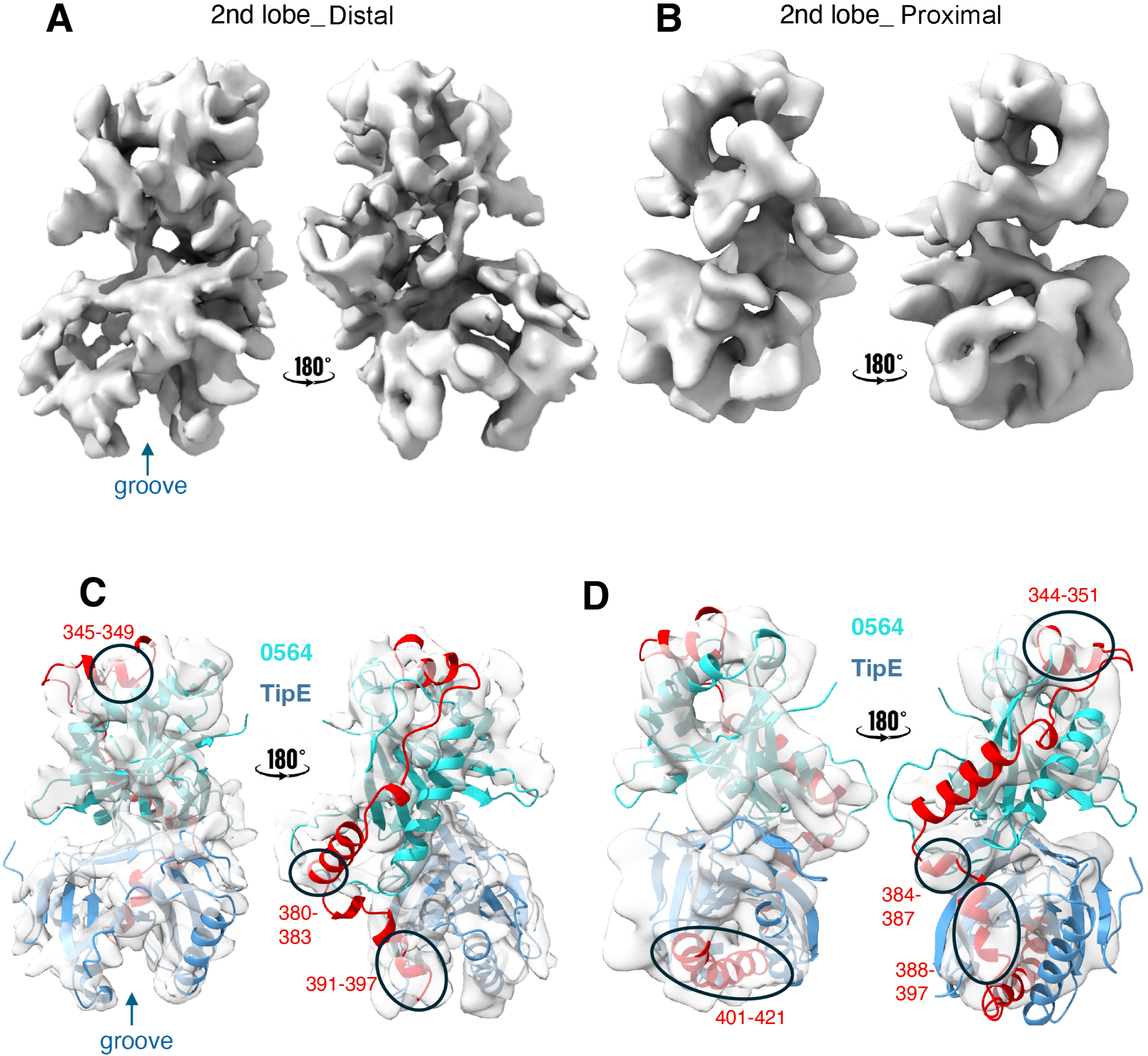
The second lobe of TelE-complex has two conformations. (A) and (B) Cryo-EM reconstruction of the LXG_TelE_ complex highlighting lobe 2_Distal and lobe 2_Proximal, contoured at a threshold of 0.104 and 0.111, respectively. (C) The AF3 model of the TelE_343-397_-0564-0565 trimer was fit to the lobe 2_Distal cryo-EM density (cross-correlation: 0.61). (D) The AF3 model of the TelE_344-421_-0564-0565 trimer was fit to the lobe 2_Proximal cryo-EM density (cross-correlation: 0.52).

A ‘third lobe’ was observed at one side of the stalk, oriented perpendicular to it (3D model on **Figure 3B**, bottom). Although the size and shape of this density might be consistent with subunit 63 being associated to the C-terminus of TelE, this assignment remains uncertain due to ambiguous 2D classification and low-resolution models. Future studies should aim to resolve the structure of 63 bound to TelE and to elucidate its binding dynamics.

Overall, our results show that the TelE-complex is organized around a stalk formed by the N-terminal region of TelE, which extends as a flexible filament linking a 61 homodimer (first lobe), a 64–65 (TipE) heterodimer (second lobe), and potentially subunit 63 (putative third lobe). Furthermore, we demonstrate that Gallo_0561 is critical for stabilizing TelE.

### The LXG-complex associates to the T7SSb pseudokinase

In a previous study we showed that the LXG domain of the T7SSb effector YxiD of *Bacillus subtilis* associates with the membrane pseudokinase YukC (an EssB homolog), a central component of the T7SSb machinery (Tassinari et al. 2022). To investigate whether a similar interaction occurs between TelE and EssB from *S. gallolyticus*, we expressed EssB in *E. coli* and purified it by affinity chromatography using an N-terminal Strep-tag (**Figure S6E**). As observed with its B. *subtilis* homolog, EssB remained stable without detergent after exchange with amphipols. We took advantage of this detergent-free condition to assess its interaction with the TelE-complex, thereby avoiding potential dissociation of the polytopic membrane protein 63 from the complex. Purified TelE-complex and EssB were mixed in an equimolar ratio and subjected to small-scale affinity purification targeting the His-tag of subunit 59 (LapE1). The eluted sample was analyzed by Western blot using anti-Strep tag to detect EssB in parallel with antibodies raised against the TelE-complex (**Figure 5A**). These results demonstrate that a stable interaction occurs between the TelE-complex and the T7SSb’s component EssB *in vitro*. Overall, our results support a model where the stalk of the TelE-complex mediates its interaction with both the pseudokinase and ATPase subunits of T7SSb, possibly causing the dissociation of the proteins bound to the middle and C-terminal domains of TelE (**Figure 5B**).

**Figure 5:**
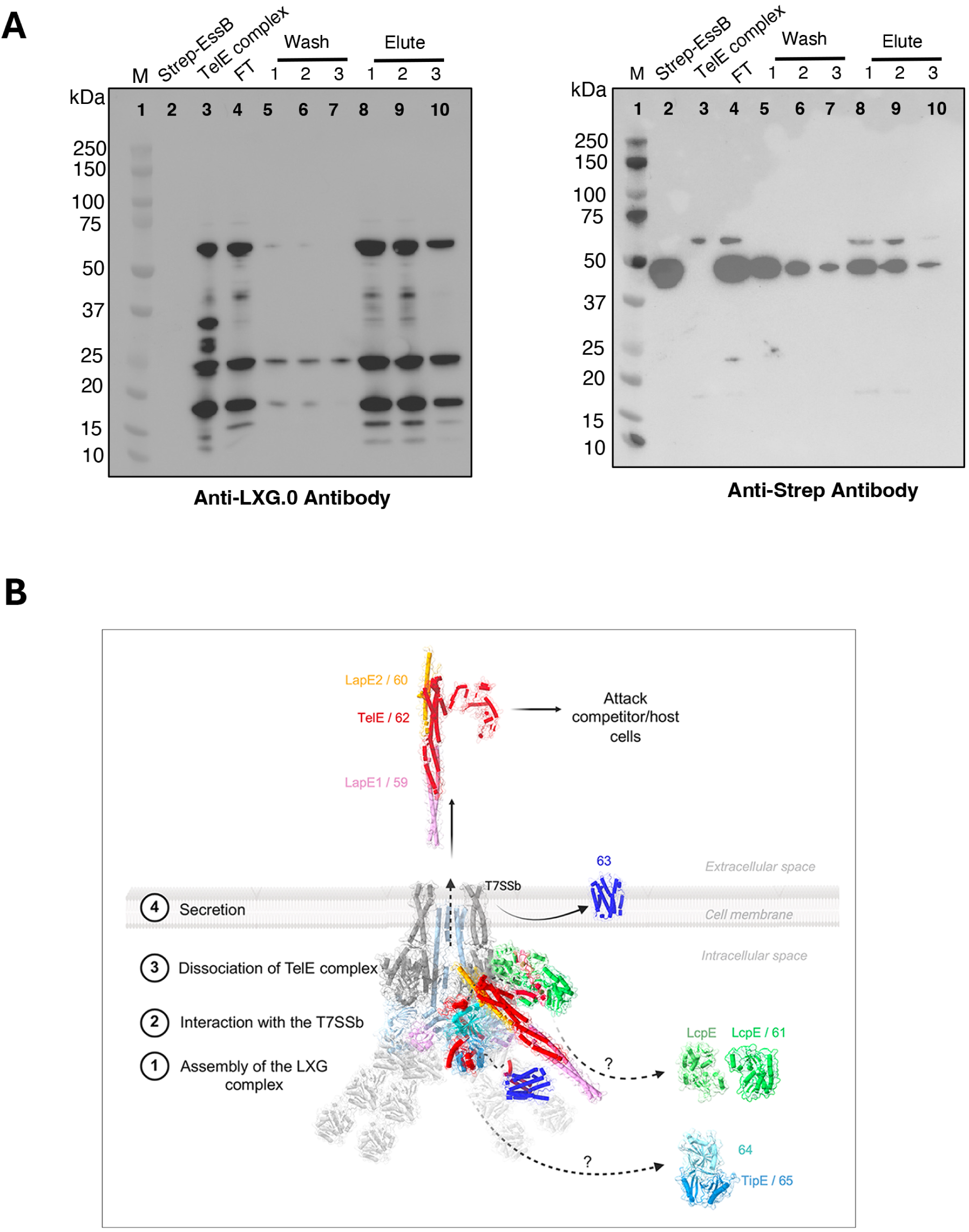
The TelE-complex associates with the pseudokinase subunit of T7SSb. (A) Cellular lysate containing His-tagged TelE-complex and Strep-tagged EssB loaded onto a NiNTA column was washed sequentially with buffers containing 5 mM (wash 1), 20 mM (wash 2), and 50 mM imidazole (wash 3), and eluted in multiple fractions with 250 mM imidazole (elute 1, 2 and 3). Samples from each steps were analyzed using a 4-20% polyacrylamide SDS-PAGE and Western blot, using antibodies raised against the LXG_TelE_-complex (left) and the Strep-tag (right). Purified TelE-complex and EssB were loaded as a positive control for anti-LXG_TelE_ and anti-Strep, respectively. (B) Proposed model of the interaction between the TelE-complex and the T7SSb machinery. The T7SSb machinery proteins EsaB/EssB/EssC ATPase has been described earlier to assemble in a 2:4:2 ratio (Oka et al. 2025). The assembled complex predicted using AlphaFold 3 is shown in grey. In the model three steps are visualized: 1) interaction of the stalk of the TelE-complex interacts with EssB and the ATPase of the T7SSb machinery; 2) dissociation of the cytosolic and membrane subunits from the TelE-complex; 3) translocation of the 59-60-62 trimer through the T7SSb machinery.

## DISCUSSION

In this study, we present the complete architecture of a seven-protein LXG complex prior to secretion via the T7SSb machinery. Previous studies characterized different LXG effectors and associated proteins in partial assemblies. Our analysis shares similarities with results on homologous subcomplexes, while revealing novel features that extend current models of LXG assemblies (Yang et al. 2023, Klein et al. 2024, Alcock et al. 2025). To our knowledge, this is the first report of a fully assembled LXG-complex, organized into a physiologically relevant model comprising three flexible domains (“lobes”) connected to a long narrow shaft (“stalk”).

We first showed that the α-helical proteins Gallo_0559 and Gallo_0560 (renamed LapE1 and LapE2) are required for TelE secretion, consistent with previous reports in *Streptococcus intermedius* and *Staphylococcus aureus* (Klein et al. 2022, 2024 and Yang et al. 2023). Our X-ray structure shows that these proteins interact with the LXG domain of TelE to form the LapE1-LapE2-TelE_1-215_ complex (PDB 9TMG), similarly to the *S. intermedius* LapA3-LapA4-TelA_LXG_ complex (PDB 8GMH) (RMSD of 1.06 with 182 Cα atoms aligned) (Klein et al. 2024). Despite this structural similarity, the Lap proteins associated with the TelE- and TelA-complexes belong to different DUF families (DUFs 5082-3130 *vs* DUFs 5082-5344). Since more structures are becoming available guiding reliable AF3 models prediction, a structural classification may prove more informative than a DUF-based annotation.

Another structural similarity between the crystallographic structure presented here and previous models is given by the two central antiparallel β-strands that interact with each Lap protein and may have a conserved, functional role (**Figure S4C**).

LXG-binding proteins also include DUF4176 proteins, a family of LXG-specific molecular chaperones required for LXG effector stability and facilitating its export (Gkragkopoulou et al. 2025, Wu et al. 2025). In *SGG* strain UCN34, Gallo_0561 -also a DUF4176 protein-acts as a TelE specific chaperone. Its absence abrogated TelE secretion, without affecting the secretion of other two LXG proteins homologous to TelC (Gallo_1068 and 1574, **Table S1**). We further showed that Gallo_0561 stabilizes the ‘middle’ and the C-terminal regions of TelE. Thus, we propose to rename Gallo_0561 as LcpE (LXG chaperone of TelE) to follow the nomenclature proposed in Gkragkopoulou et al. 2025. Our structural analysis shows that LcpE interacts with TelE in a region predicted to be mainly α-helical, with no disordered segments between residues 220 and 270 according to PSIPRED (McGuffin et al. 2000) or AF3 (Abramson et al. 2024).

We showed previously that TelE’s pore-forming activity is neutralized by the immunity protein TipE (a member of the DUF5085 family) when overexpressed in *E. coli* (Teh et al. 2023). EsxX, a TelE homolog, recently characterized in *S. aureus* was shown to depolarize cell membrane from both cytosolic and extracellular compartments (Alcock et al. 2025). In *S. aureus*, a pair of polytopic membrane proteins (ExiCD) was shown to protect against extracellular EsxX and a pair of cytosolic proteins (ExiAB was shown to protect against cytoplasmic EsxX (Alcock et al. 2025). Similarly, TelE can be toxic from the cytosolic compartment (Teh et al. 2023) as well as from the external compartment (**Figure 2C**). Strikingly, the gene *gallo_0563* encoding a polytopic membrane protein is located immediately downstream TelE. While it is tempting to speculate that Gallo_0563 is a ExiCD homolog conferring a similar protective mechanism, we showed a main difference in that Gallo_0563 associates to form the soluble TelE-complex. If Gallo_0563 indeed functions as TelE immunity protein, it is intriguing to understand how the translocation of this membrane protein to the bacterial membrane is coordinated during TelE secretion via the T7SSb machinery.

Cryo-EM analysis of this seven-protein complex was highly challenging due to the difficulties posed by the sample behavior (air-water interface instability, preferential particle orientation, local clustering) combined with the analysis of its flexible appendages. Fitting to AF3 models, the map of the stalk linked to the first lobe showed that the latter is constituted by a dimer of LcpE (61) interacting with TelE G222-Ile267. This fragment is homolog to the ‘middle region’ previously described for TelA and TelB (**Figure S7**) (Gkragkopoulou et al. 2025). The models presented here for the LcpE dimer is highly similar to the one for LtcA and LtcB from *S. intermedius*, except for the interaction with the effector. Here we propose that the LXG effector binds two LcpE proteins, in contrasts with the 1:1 ratio shown for *S. intermedius*. Interestingly, Gkragkopoulou et al. also obtained a crystalline form with a 2:1 ratio but considered it non-physiological.

### Interaction with T7SS and conclusions

Based on previous evidence and these results, we propose a revised model for the interaction between LXG complexes and the T7SSb secretion machinery (**Figure 5B**). In this model, binding of LXG complexes on the T7SSb machinery is mediated by conserved features located on the stalk, possibly including the central β-strands observed in different LXG-systems. The presence of two conserved motifs in LapE2, similar to those found in T7SSa substrates (Klein et al. 2024; Alcock et al. 2025), suggested that the stalk contacts the ATPase. We propose that an additional contact occurs between the LXG complex and the pseudokinase (EssB), as illustrated in the model proposed in **Figure 5B**. This is in accordance with previous data showing interaction between the LXG-domain and the membrane pseudokinase of the T7SSb in *B. subtilis* (Tassinari et al. 2022). Here, we observed that not only the LXG-effector, but all the pre-secretion complex, binds stably to this T7SSb subunit. Recently, the first model of a T7SSb from *B. subtilis* showed that the pseudokinase domains are entangled with the FHA domains of the ATPases in the cytoplasmic side (Oka et al. 2025). It is therefore possible that LXG-complexes contacts the pseudokinase subunit and the nearby ATPase concomitantly, initiating its translocation through the secretion machine.

We also propose that the LXG-complexes organize in a modular way with associated proteins binding the toxic effector in structurally independent sub-domains consisting of independent lobes attached to the conserved stalk. This modularity might facilitate TelE transport through the T7SS without decreasing the efficiency in detoxification. Proteins associated in the lobes to the LXG effector, need to dissociate before entering the T7SSb pore formed by the hexameric ATPases (Oka et al. 2025). This agrees with the literature on homolog systems and with our secretome results showing that only the Lap proteins, LapE1 and LapE2, are co-secreted with TelE. The fact that the membrane protein Gallo_0563 is also associated to the pre-secretion complex, but not found in the secretome, suggests that it transitions to the membrane during the complex’ secretion. It is tempting to speculate that the modular organization observed here, coupled to a limited surface of interaction between the effector C-terminus and the cytosolic subunits, might be instrumental to their dissociation without delaying the secretion.

## MATERIAL AND METHODS

### Cultures, bacterial strains, plasmids and oligonucleotides

*Streptococcus gallolyticus subsp. gallolyticus* (*SGG*) strain UCN34 (Rusniok et al. 2010) and its derivatives were grown at 37°C in Todd-Hewitt broth alone (TH) or supplemented with yeast extract 0.5% (THY) in standing flasks or agar plates. When appropriate, 10 µg/mL of erythromycin was added for plasmid maintenance.

### Construction of marker less deletion mutants in *SGG* UCN34

In-frame deletion mutants were constructed as described previously (Danne et al. 2013). Briefly, the 5’ and 3’ flanking the region to delete were amplified and assembled by overlap extension PCR. The PCR products were subsequently cloned into the thermosensitive shuttle vector pG1. Once transformed in UCN34, the bacteria were cultured at 38°C with erythromycin to select for the cells containing the plasmid integrated into the chromosome by homologous recombination. About 4 single crossover integrants were serially passaged at 30°C without antibiotic to facilitate the second event of homologous recombination, to obtain in-frame deletion mutant or reversion to the WT strains. Mutant strains harboring in-frame deletions were screened by PCR and confirmed by sequencing the chromosomal DNA flanking the target region. All the strains and primers used in this study are listed in **Table 1**.

**Table 1:**
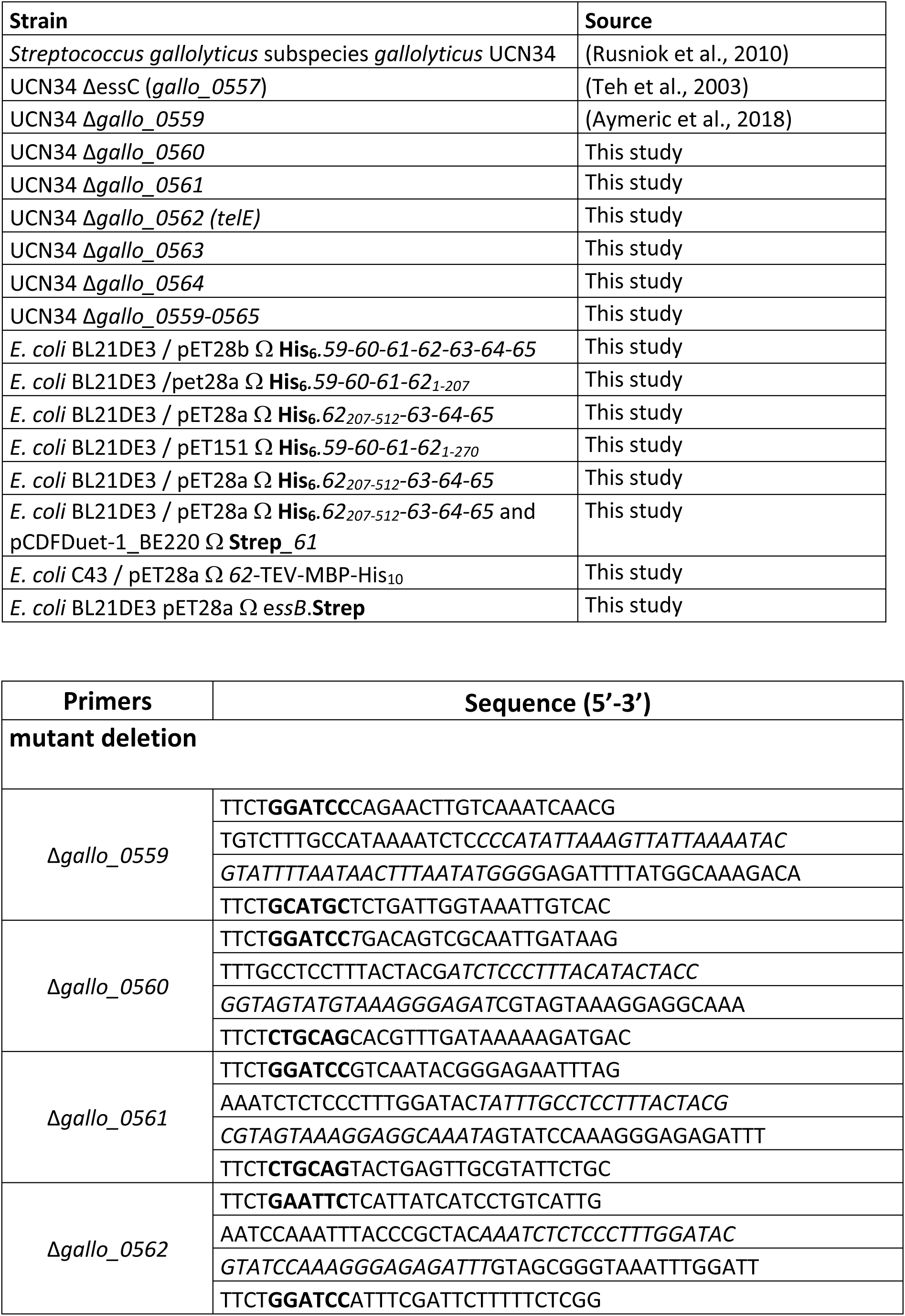
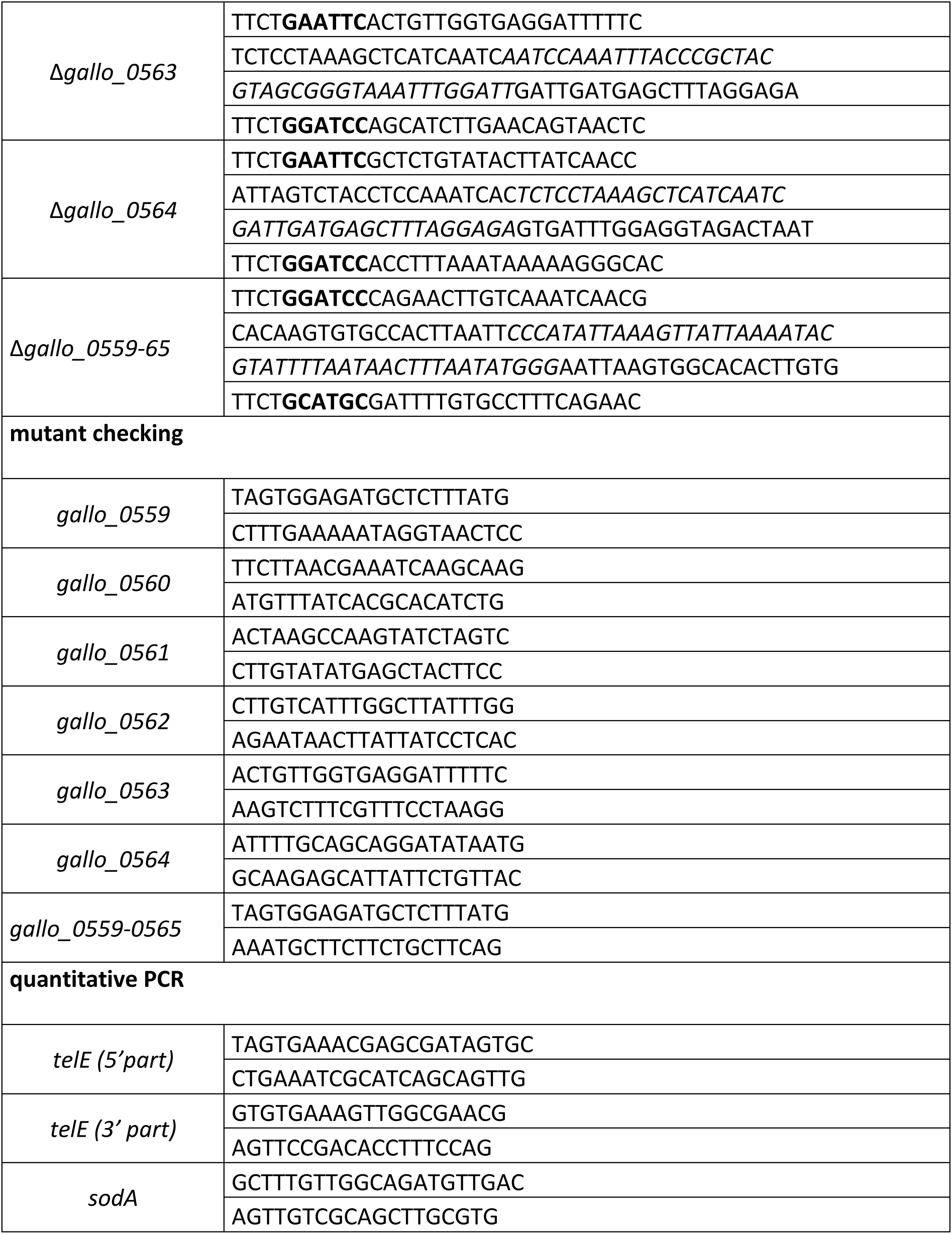
Strains and primers used in this study.

### Growth curves of *SGG* strains

Bacteria from −80°C collection were plated on Todd-Hewitt plates and incubated overnight at 37°C. The next day, bacteria were resuspended in PBS and adjusted to get an optical density measured at 600 nm, OD_600_ of 2 per mL. Next, each bacterial culture was diluted in TH broth (50 µl in 950 µl of TH) to OD_600_ of 0.04. A total of 200 µL of this suspension was dispensed into sterile 96-well plates in triplicate. The culture was incubated at 37°C without shaking on Multiskan Go plate reader device (Thermo Scientific). The OD_600_ was measured every 15 min and with a short mix before the reading.

### Flow cytometry

*E. coli* GM48 cells grown to exponential phase (OD_600_ = 0.5), were first washed and resuspended in PBS. The cells were subsequently incubated with CFSE for 20 min at 37°C. Then Hoechst 33342 (1/1000) was added to the sample for 5 min to visualize bacterial DNA. Following incubation, the stained cells were washed and resupended in PBS. Stained cells in 100 µL aliquots were incubated with detergents, control buffer or purified TelE at 37°C for 10 min under gentle agitation. The samples were acquired on a MACSQuant YGV Analyzer apparatus (Miltenyi Biotec) and data were analyzed using FlowJo X software.

### Quantitative RT-PCR

Bacterial colonies pre-cultured overnight on TH agar plate at 37°C were diluted to OD_600_ of 0.05 in 25 mL TH and grown at 37°C until late exponential growth phase (OD_600_ of 0.8-1), when the expression of T7SSb genes is maximal. A total of 20 mL of the bacterial culture concentrated by centrifugation 4°C, 7000 g for 10 min was subjected to total RNA extraction (FastRNA PRO™ BLUE Kit, MP Biomedicals). Residual DNA from the RNA samples was removed using the TURBO™ DNase kit (2U/μl, Ambion™ / Thermo Fischer Scientific). RNA quality was assessed by agarose gel electrophoresis, whereas the RNA concentrations were measured using Nanodrop ND-100 spectrophotometer (ThermoFischer). When necessary, RNA extracts were stored at −80°C. Total RNA (500ng) was reverse transcribed into cDNA using random hexamer oligonucleotides with Superscript IV reverse transcriptase (Invitrogen, ThermoFisher Scientific). Quantitative real-time PCR was performed in 15 µL mix containing 7.5 µL Power SYBR Green PCR Master Mix (Applied Biosystems), 0.75 µL gene-specific primers (10 mM) and 5 µL of a 100-fold diluted cDNA on a LightCycler 480 Real-Time PCR System (Roche). Relative gene expression levels were calculated using the comparative 2^−ΔΔCt^ method (Livak and Schmittgen 2001). *SodA* was used as the reference gene for normalization. Each assay was performed in triplicate with three independent cultures.

### Secretome analyses

To assay for protein secretion, 2 mL of bacterial cultures grown in M9 medium supplemented with 0.5% of yeast extract and 0.5% glucose were grown at 37°C until OD_600_ of 0.8-1 and harvested though centrifugation at 4,000 g for 5 min at 4°C. The supernatant was filtered through 0.1 um filter and proteins were precipitated with TCA 10% overnight at 4°C. Precipitated proteins were then recovered by centrifugation at 13,000 g for 30 min, washed with acetone, dried and stored at −20 °C until further analysis. Proteins were digested using an adapted S-Trap™ protocol (Zougman et al. 2014) with S-Trap mini columns (ProtiFi™). Samples were solubilized in SDS lysis buffer (5% SDS, 50 mM TEAB, pH 7.55), reduced with TCEP, and alkylated with iodoacetamide. Proteins were acidified, diluted in S-Trap binding buffer (100 mM ammonium bicarbonate in 90% methanol), and loaded onto S-Trap columns. After washing, proteins were digested on-column overnight at 37 °C with sequencing-grade trypsin. Peptides were sequentially eluted, pooled, dried by vacuum centrifugation, and stored at −20 °C until LC–MS/MS analysis.

Peptides were resuspended in 0.1 % formic acid and analyzed using an Orbitrap Q Exactive Plus mass spectrometer (Thermo Fisher Scientific) coupled to an EASY-nLC 1200 system. Peptides were separated on a 75 µm inner diameter reversed-phase column with a linear acetonitrile gradient in 0.1 % formic acid at 250 nL/min. Data were acquired in data-dependent acquisition mode with full MS scans at 70,000 resolutions followed by MS/MS scans of the top 10 most intense precursors using HCD fragmentation.

Raw data were processed using MaxQuant v2.1.4.0 with the Andromeda search engine (Cox et al. 2011). Spectra were searched against the *Streptococcus gallolyticus* subsp. *gallolyticus* UCN34 proteome (2,217 entries,17.04.2024). Carbamidomethylation of cysteine was set as a fixed modification, and methionine oxidation and protein N-terminal acetylation as variable modifications. Trypsin specificity with up to two missed cleavages was allowed. Peptide and protein false discovery rates were controlled at 1%. The proteomics data have been deposited in the ProteomeXchange Consortium via PRIDE under accession PXD075233.

### Purification of the TelE -complex and sub-complexes

*Escherichia coli* BL21(DE3) cells were transformed with the following plasmids for TelE-complex and sub-complexes expression: (i) pET28bΩHis_6_*.0559-0560-0561-0562-0563-0564-0565* (LXG.0 Construct), (ii) pET28aΩhis_6_*.59-60-61-62_1-207_* (LXG.1N construct), (iii) pET28aΩ*his_6_.62_207-512_-63-64-65* (LXG.2 construct), or (iv) pET28aΩhis_6_*.62_207-512_-63-64-65* (LXG.1C construct) along with pCDFDuet-1_BE220/Strep*_0561*. The various gene constructs and cloning were ordered at GenArt Services (ThermoScientific). When appropriate, antibiotics were supplied at 50 μg/mL for kanamycin, 100 μg/mL for ampicillin and 50 μg/mL for spectinomycin. Cells were incubated at 37 °C overnight for growth.

Start cultures was prepared from single colonies inoculated into LB and incubated at 37°C while shaking for 2 h. The start culture was subsequently diluted 1:50 into fresh LB medium containing the corresponding antibiotics. Cultures were incubated at 37°C to OD_600_ of 0.8. Protein expression was induced with 0.5 mM of isopropyl b-D-1-thiogalactopyranoside (IPTG), followed by incubation for 3 h at 37°C with shaking. Cells were harvested by centrifugation at 3,300 g for 20 min at 4°C and resuspended in cold lysis buffer A (50 mM Tris-HCl pH 8.0, 250 mM NaCl, one tablet of protease inhibitor cocktail; Roche). Benzonase (25 U/mL) and lysozyme (0.1 mg/mL) were added, and cells were lysed by sonication (10-30 s on/off cycles for 10 min at 40% amplitude; Sonics VibraCell, Fischer Brand). Cell debris was removed by centrifugation at 8,000 g for 10 min at 4°C. For constructs containing membrane-associated components, lysates were further ultracentrifuged (Beckman Coulter XPN100) at 100,000 g for 30 min at 4 °C to separate soluble and membrane fractions. Membrane pellets were resuspended in buffer A and solubilized in the presence of 1 % (w/v) n-dodecyl-β-D-maltoside (β-DDM). Both soluble and solubilized membrane fractions were processed for affinity purification.

Imidazole was added to a final concentration of 5 mM prior to loading onto Ni-NTA resin (Cytiva), either using a 5 mL column (for full-length LXG_TelE_) or 200 µL resin (for subcomplexes), pre-equilibrated with buffer A containing 5 mM imidazole. Binding was performed at 4°C (overnight for small-scale purifications). For full-length LXG_TelE_, a gradient from 5 mM to 50 mM imidazole in Buffer A was used for washing at 2 mL/min for 20 min. Bound proteins were eluted in Buffer A containing varying levels of imidazole ranging from 50 mM to 250 mM imidazole, at a flow rate of 2mL/min for 30 min. For LXG_TelE_ subcomplexes, resins were washed with 10 column volume ‘CV’ of buffer A followed by buffer A supplemented with 50 mM imidazole. Proteins were eluted in Buffer A containing 120 and 250 mM imidazole. Eluted proteins were collected in 1-mL fractions (for full-length LXG_TelE_) and 200-µL fractions (for subcomplexes). These eluted fractions were analyzed by SDS-PAGE, and the protein identity was confirmed by mass spectrometry on the peptides recovered from the excised gel bands. For stability assays, purified TelE complex was adjusted to 0.5 mg/mL (approximately 2.5 µM) and concentrated using a 100 kDa MWCO centrifugal filter (Vivaspin).

### Purification of TelE-MBP-His_10_

*E. coli* C43 cells transformed with pET28aΩ*0562*-TEV-MBP-His_10_ was first plated on LB agar supplemented with Kan 50 μg/mL and 0.5% glucose. For the starter culture, a single colony of transformed cells was inoculated in 100 mL of LB supplemented with Kan 50 μg/mL and 0.5 % glucose and grown to an OD_600_ of 0.7 (about 3 h at 37 °C, while shakin). The cells collected by centrifugation at 4000 g for 10 min were diluted into 4 L of TB medium supplemented with Kan (50 *μ*g /mL). Cells were incubated at 37 °C until OD_600_ of 0.8. Then IPTG was added to a final concentration of 0.5 mM, and the bacterial culture was incubated for 2 h at 30°C shaking. Bacterial culture was spiked with 1 mM Phenylmethanesulfonyl Fluoride (PMSF) and cells were harvested by centrifugation at 3,300 g for 15 min at 4°C. The resulting cell pellet was flash-frozen in liquid nitrogen and stored at −80°C until protein purification. During purification, the sample was kept on ice unless otherwise specified. To purify TelE, cells were resuspended in lysis buffer (50 mM HEPES pH 7.5, 50 mM NaCl, 1 mM PMSF) and lysed by sonication (10s on-off pulse cycles for 10 min, at 40% amplitude; Sonics Vibracell, Fischer Brand). Unbroken cells were removed by centrifugation at 8,000 g for 10 min at 4 °C. The resulting supernatant was ultracentrifuged (Beckman Coulter XPN100) at 100,000 g for 45 min at 4°C to separate the soluble and membrane proteins. Pelleted membrane proteins were resuspended to a final volume of 60 mL in Buffer B (50 mM HEPES pH 7.5, 150 mM NaCl, 1mM PMSF and 1 tablet of protease inhibitors; Roche). Solubilization was achieved by adding β-DDM to a final concentration of 1% final achieved with stirring for 30 min at 4 °C. The solubilized membrane proteins were subsequently loaded at 2 mL/min on to the pre-equilibrated 5 mL Mannose binding protein (MBP) column (Cytiva) in Buffer C (50 mM HEPES pH 7.5, 150 mM NaCl, 1 mM TCEP, and 0.02% β-DDM) mounted on an Akta purifier (GE HealthCare). Protein-bound MBP resins were washed with 50 mL of Buffer C and TelE-MBP-His_10_ was eluted using the same buffer supplemented with 10 mM maltose (sigma). The eluted fractions were analyzed by SDS-PAGE stained by Coomassie followed by Western blotting. TelE-MBP-His_10_ was then treated with TEV protease at a 1:15 (w/w) in the presence of 1 mM DTT. The reaction was incubated overnight at 4°C under gentle rotation. Cleaved products as well as TEV protease were then removed by passing the sample through Ni-NTA agarose resin (Qiagen, Germany) equilibrated in Buffer B (see above). The flow-through from Ni-NTA was collected after 2 h of incubation. Samples were evaluated by SDS-PAGE stained by Coomassie. The purified TelE was then used for functional assays.

### Mass Photometry

Molecular mass measurements were performed using a OneMP mass photometer (Refeyn, Ltd.). Coverslips (No. 1.5 H, 24 × 50 mm, Marienfeld) were cleaned through three sequential washes with isopropanol and Milli-Q water, followed by drying with a stream of nitrogen. LXG_TelE_ complex was diluted to 20 nM in 50 mM Tris, pH 8.0, 250 mM NaCl. To measure the molecular mass of the individual proteins and complexes, a clean coverslip was mounted onto the mass photometer, and a reusable gasket (CultureWell™ Reusable Gasket, Grace Bio-Labs) was placed on top. Each well was filled with 18 µl of sample buffer, followed by the addition of 2 µl of the diluted sample. The adsorption of single molecules onto the glass surface was monitored for 60 s using AcquireMP software (Refeyn Ltd, Version 2.3.0). Molecular mass was calculated by converting ratiometric contrast values using a calibration curve generated with Bovine Serum Albumin (BSA) and its oligomers, covering a range from 66 to 264 kDa (monomer to tetramer). All mass photometry movies were analyzed using DiscoverMP (Refeyn Ltd, Version 2.3.0).

### LUV Permeabilization Assay

Permeabilization-induced dye leakage was assessed using large unilamellar vesicles (LUVs) encapsulating the fluorophore ANTS (8-aminonaphthalene-1,3,6-trisulfonic acid) and its quencher DPX (p-xylene-bis-pyridinium bromide). Lipid vesicles (final lipid concentration, 4 mM) were prepared from a mixture of DOPE and DOPG (75:25) containing 20 mM ANTS and 60 mM DPX by the reverse-phase evaporation method. Vesicles were subsequently extruded sequentially through 0.8-, 0.4-, and 0.2-μm polycarbonate membranes to generate LUVs, as described previously (Karst et al. 2012). Non-encapsulated probes were removed by buffer exchange using a Sephadex G-25 column equilibrated with Buffer B (20 mM HEPES, 150 mM NaCl, pH 7.4). The resulting LUV suspensions were monodisperse, with a mean hydrodynamic diameter of approximately 180 nm, as determined by dynamic light scattering using a NanoZS instrument (Malvern) (Subrini et al. 2013). For permeabilization assays, LUVs were incubated at 25 °C, and ANTS fluorescence was monitored continuously (λ_ex_ = 360 nm, λ_em_ = 515 nm, bandwidths 5 nm) to set a stable baseline. Proteins were then added to the vesicle suspension, and changes in fluorescence intensity were recorded over time.

### Crystallization

Limited proteolysis was performed using Trypsin in a 1:50 (w/w) ratio with the TelE-complex, as measured by Nanodrop at 280 nm, and the digestion was carried out on ice. After 1 h, EDTA-free Protease Inhibitors (Roche) were added (final concentration 10x) and the mixture injected on a SEC Superdex 200 Increase 10/300 GL column (Cytiva). Three main elution peaks were obtained (**Figure S3A**). Fractions corresponding to each peak were pulled and applied on a mPAGE™ 4-20% Bis-Tris Precast gel for SDS-PAGE followed by Coomassie staining. Peaks 1, 2 and 3 were concentrated on 50 kDa MWCO filters, reaching final concentrations of around 3.0, 4.0 and 4.5 mg/ml, respectively.

Crystallization screenings were performed on hanging drop plates at 4 °C, by mixing equal volumes of samples and crystallization solutions (0.4 μL drops); crystals from ‘Peak #3’ sample were observed in the presence of 0.2 M CaCl2, 0.1 M Na Acetate pH 4.5, 30% MPD. Mass spectrometry of the SDS-PAGE gel bands from ‘Peak #3’ identified fragments of subunits 59, 60, 61, 62 and 64. Crystals were flash-frozen in liquid nitrogen and a diffraction dataset collected at the ESRF synchrotron. Data integration and scaling were performed with autoPROC (Vonrhein et al., 2011), applying anisotropic scaling via STARANISO. The structure was solved by molecular replacement through the program PHASER (McCoy et al. 2007), using a generated Alphafold 3 model (Jumper et al. 2021) comprising the N-terminal domain of the pore-forming toxin TelE and two Lap subunits, identified in the sample. Model rebuilding in electron density maps was performed with COOT (Emsley et al. 2010). Reciprocal space refinement was carried out with BUSTER, applying local structure similarity restraints for non-crystallography symmetry (NCS) (Smart et al. 2012)and a Translation-Libration-Screw (TLS) model, defined as one TLS group per polypeptide chain. Model validation was performed with MOLPROBITY (Williams et al., 2018) and the validation tools in PHENIX (Liebschner et al., 2019). Figures were generated with PyMol (Schrödinger, LLC) and ChimeraX (Meng et al. 2023). Data collection, refinement and model quality statistics are reported in **Table S2**. Multiple Sequence Alignment was performed using Clustal Omega (Madeira et al. 2024) on 10 different LXG-proteins and visualized with WebLogo software (Crooks et al. 2004).

### CryoEM sample preparation, data collection and structure determination

CHAPSO was added to the sample before freezing, to protect the protein from air-water interface damages. Vitrification was carried out using a Vitrobot plunger (Thermo Fischer Scientific). 3 μl of the TelE-complex at 0.1 mg/mL was applied to glow discharged Aufoil 1.2/1.3 300 mesh grids (Quantifoil), blotted for 5 s, blot force 0, at 8 °C, 100% humidity, and plunge-frozen in liquid ethane. Data were collected in a 200 kV Glacios electron microscope (Thermo Fisher Scientific) equipped with a Falcon 4i Direct Detection camera (Gatan) at a total dose of 40 e-/Å^2^ at the specimen level and a sampling of 0.57 Å/pixel. A total of 8,356 movies were collected. Single Particle Analysis was performed using CryoSPARC (Punjani et al., 2017). Particles were manually selected to create an initial template for subsequent template-based picking. Movies were subjected to patch-based motion correction, contrast transfer function (CTF) estimation and manually curated to 8,356 micrographs for downstream processing. The detailed workflow is illustrated in **Figure S8**. Denoised micrographs were used for processing of Lobe 2. Initial manual picking of 475 particles from a subset of 23 micrographs was used to generate 5 templates for subsequent automated particle picking. **Analysis of Stalk + lobe 1:** after multiple rounds of template picking and 2D classifications cleaning we selected 1,166,264 that were used for Ab initio reconstruction into five classes. Heterogeneous refinement with five classes and non-uniform refinement of (Class 0), yielded a reconstruction at 6.1 Å resolution. The particles were further subjected to 3D classification into 10 classes and from which 6 classes were picked for additional rounds of 2D classification, resulting in a curated dataset of 154,037 particles. Amongst these, 65,521 particles were selected for focused refinement. These particles were used for Ab initio reconstruction into one class, followed by non-uniform refinement, producing a reconstruction corresponding to the stalk with one lobe at a final resolution of 4.5 Å. **Analysis of the Stalk + 3 lobes**: a subset of 111,904 particles was selected for Ab initio reconstruction into one class, followed by heterogeneous and local refinement, yielding a reconstruction including 55,733 particles at 6.5 Å resolution. **Analysis of the Second Lobe in the Distal conformation**: 42,993 particles were subjected to Ab initio reconstruction into one class, followed by non-uniform and local refinement, yielding a reconstruction at 4.2 Å resolution. **Analysis of the Lobe 2 in the Proximal conformation**: 19,546 particles were subjected to Ab initio reconstruction into one class, followed by non-uniform and local refinement, yielding a reconstruction at 5.2 Å resolution. Interpretation of 3D maps obtained by cryo-EM was performed using AlphaFold 3 models (Abramson et al. 2024).

### Purification of EssB

A single colony of *E. coli* BL21(DE3) transformed with pET28a Ω strep.*EssB* was inoculated into 100 mL of LB supplemented with Kan 50 µg/mL. After overnight growth at 37°C with shaking, the seed culture was diluted 1:50 in 2 L of LB medium supplemented with Kan 50 µg/mL and incubated at 37°C with shaking to an OD_600_ of 0.8. Expression was induced by the addition of 0.5 mM IPTG and incubated overnight at 16 °C while shaking. Cells were collected by centrifugation at 6,000 g for 20 min at 4°C. During purification, the sample was kept on ice unless otherwise specified. Cells were resuspended in 100 mL of cold lysis buffer (50 mM HEPES pH 8, 175 mM NaCl, 5% glycerol, 1 mM EDTA, and 2 tablets of protease inhibitors; Roche) and then incubated at room temperature (RT) for 20 min under moderate shaking in the presence of 0.5 mg/mL lysozyme (Sigma-Aldrich) and for an additional 10 min after the addition of 0.1 mg/mL DNase, 5 mM MgCl_2_. Cells were lysed by sonication (10s on-off pulse cycles for 10 min, at 50 % amplitude; Sonics Vibracell, Fischer Brand). Unbroken cells were centrifuged for 15 min at 8,000 g. Membrane proteins were pelleted by ultracentrifugation using a Ti45 rotor (Beckman Coulter) running at 100,000 g for 1 h (at 4°C) and resuspended in 20 mL of cold solubilization buffer (25 mM HEPES pH 8, 175 mM NaCl, 2.5% glycerol, 1 tablet of protease inhibitors). 20 mL of 2% Cymal-6 (Anatrace) solution was added to the membrane suspension, and the mixture was incubated at RT for 30 min under moderate agitation. Insoluble material was removed by ultracentrifugation using a Ti45 rotor (Beckman Coulter) at 100,000 g for 30 min. The cleared solubilized material was loaded at 0.8 mL/min on a preequilibrated 5-mL StrepTrap column (GE Healthcare) mounted on an Akta purifier (GE healthcare) preequilibrated with purification buffer (25 mM HEPES, pH 8, 175 mM NaCl, 0.04 % Cymal-6). The affinity column was washed with 50 mL of purification buffer, and Strep-EssB was eluted using the same buffer supplemented with 2.5 mM desthiobiotin (Sigma). The eluted fractions and the intermediate purification steps were evaluated by SDS-PAGE.

### Interaction of LXG_TelE_-complex with the pseudokinase EssB

This protocol was performed between 4 and 6 °C unless otherwise stated. Amphipols A8-35 (Anatrace) was added to the purified Strep.EssB in a 1:1 ratio (w/w) in 25mM HEPES pH 8.0, 175 mM NaCl, 0.04 % cymal-6, 2.5 mM desthiobiotin. After incubation on ice for 30 min, the NaCl concentration was decreased to 90 mM and imidazole was added to 5 mM, final concentration. The LXG_TelE_ complex was added at a 1:1 molar ratio with Strep.EssB. The mixture was added to pre-equilibrated Ni-NTA resin (Qiagen), incubated in cold room under gentle rotation for 30 min. After centrifugation at 500 g for 30 s, the supernatant was removed and the resin was washed in 5 column volumes of 50 mM HEPES pH 7.5, 50 mM NaCl, 5 mM imidazole. After two additional wash steps, the bound proteins were eluted using buffer with increasing imidazole concentrations: 20 mM, 50 mM and 250 mM. The eluted proteins were loaded onto two SDS-PAGE gels for concurrent Western Blot analysis using two different antibodies against i) the LXG_TelE_-complex and ii) the Strep-tag (for Strep.EssB).

## ACKNOWLEDGEMENTS

This paper was supported by a PTR grant from Institut Pasteur to S.D., F.G. and A.C. (PTR 2023 n°708). A.G. and J.G. benefitted of the ERASMUS+ program from the European Union. We thank Auriane de Malherbe for help in RT-qPCR experiments. We acknowledge the following platforms at the Institut Pasteur: the Nanocore facility, for cryo-EM screening and data collection; the Crystallography facility, for screening and help with the crystallization screenings of the stalk; the Biophysics facility, for support with Mass Photometry. We are thankful to the ESRF (Grenoble, France) and SOLEIL (Saint-Aubin, France) synchrotron sources for granting access to their respective facilities. We also thank the following people for their precious inputs: Rafael Navaza for his help with Cryosparc set up; Rémi Fronzes for his support on the SPA analysis of the stalk-first lobe; Heddy Soufari, Pierre Legrand and all the Polaris support staff at the SOLEIL synchrotron.

